# Basolateral amygdala population coding of a cued reward seeking state depends on orbitofrontal cortex

**DOI:** 10.1101/2023.12.31.573789

**Authors:** David J. Ottenheimer, Katherine R. Vitale, Frederic Ambroggi, Patricia H. Janak, Benjamin T. Saunders

## Abstract

Basolateral amygdala (BLA) neuronal responses to conditioned stimuli are closely linked to the expression of conditioned behavior. An area of increasing interest is how the dynamics of BLA neurons relate to evolving behavior. Here, we recorded the activity of individual BLA neurons across the acquisition and extinction of conditioned reward seeking and employed population-level analyses to assess ongoing neural dynamics. We found that, with training, sustained cue-evoked activity emerged that discriminated between the CS+ and CS-and correlated with conditioned responding. This sustained population activity continued until reward receipt and rapidly extinguished along with conditioned behavior during extinction. To assess the contribution of orbitofrontal cortex (OFC), a major reciprocal partner to BLA, to this component of BLA neural activity, we inactivated OFC while recording in BLA and found blunted sustained cue-evoked activity in BLA that accompanied reduced reward seeking. Optogenetic disruption of BLA activity and OFC terminals in BLA also reduced reward seeking. Our data suggest that sustained cue-driven activity in BLA, which in part depends on OFC input, underlies conditioned reward-seeking states.

## INTRODUCTION

A foundational building block of motivated behaviors, such as food seeking or threat avoidance, is Pavlovian conditioning, wherein animals develop behavioral responses to sensory cues that predict positive or negative outcomes (Pavlov and Anrep, 1927). The basolateral amygdala (BLA) is a major site of plasticity underlying this cue-outcome associative learning (McKernan and Shinnick-Gallagher, 1997; Fanselow and LeDoux, 1999; Tye et al., 2008; Clem and Huganir, 2010). An important ongoing area of investigation is how in vivo neural dynamics in the BLA direct cue-elicited motivated behaviors (Janak and Tye, 2015). In addition to considerable work on fear conditioning and the BLA, there is also a strong body of work demonstrating robust neural modulation in the BLA in response to reward-predicting cues (Muramoto et al., 1993; Schoenbaum et al., 1998; Paton et al., 2006; Belova et al., 2007; Tye and Janak, 2007; Ambroggi et al., 2008; Tye et al., 2008; Shabel and Janak, 2009; Sangha et al., 2013; Beyeler et al., 2016; Burgos-Robles et al., 2017; Lee et al., 2017; Kyriazi et al., 2018; Zhang and Li, 2018; Lutas et al., 2019). Compellingly, cue-evoked activity in the BLA changes across learning stages (Muramoto et al., 1993; Schoenbaum et al., 1998, 1999; Paton et al., 2006; Herry et al., 2008; Grewe et al., 2017; Zhang and Li, 2018; Lutas et al., 2019; Lee et al., 2021; Zhang et al., 2021). A remaining question is how within-trial neural dynamics in BLA evolve during the execution of learned cue-driven behaviors.

Here, using a combination of electrophysiological, pharmacological, and optogenetic approaches in freely-moving rats, we examined the relationship between neural activity in the BLA and ongoing behavior during the acquisition and extinction of a Pavlovian reward-seeking task. We also studied the contribution of an important input to the BLA, the orbitofrontal cortex (OFC) (Saddoris et al., 2005; Morrison et al., 2011; Lucantonio et al., 2015; Sharpe and Schoenbaum, 2016; Arguello et al., 2017), to BLA population activity during this task.

We found that, during acquisition of the task, persistent population activity in BLA increasingly discriminated between the conditioned stimuli and correlated with the acquisition of conditioned behavior. During extinction, neural and behavioral discrimination concurrently decreased. Interestingly, the persistent activity triggered by CS+ presentation terminated upon reward acquisition, and optogenetic disruption of BLA neurons during CS+ presentation impaired expression of conditioned reward seeking, suggesting that persistent BLA neural dynamics support conditioned reward seeking. Finally, inactivation of OFC reduced conditioned behavior and cue-evoked BLA dynamics, and photoinhibition of OFC input to BLA also reduced reward seeking, indicating that the influence of BLA on reward seeking is dependent on input from OFC. Overall, these findings demonstrate the importance of the BLA circuit in orchestrating ongoing conditioned behavior across learning.

## RESULTS

### Sustained CS+ activity in BLA tracks acquisition of conditioned reward seeking

To study how basolateral amygdala (BLA) neural dynamics evolve across learning, we recorded BLA single units in rats (n=18) as they underwent a Pavlovian conditioning procedure (Figure 1A-C). During the “Conditioning” phase, one 30-sec sound (CS+, tone or white noise) was repeatedly paired with sucrose delivery, and the second sound (CS-) was paired with nothing (Figure 1A). Rats performed 7 days of Conditioning and then two additional sessions in which the association was extinguished (“Extinction” and “Extinction Recall”) (Figure 1B). Rats acquired robust conditioned reward seeking across Conditioning, as evidenced in their increased time spent in the reward port during CS+ relative to CS(Figure 1D-E).

**Figure 1.**
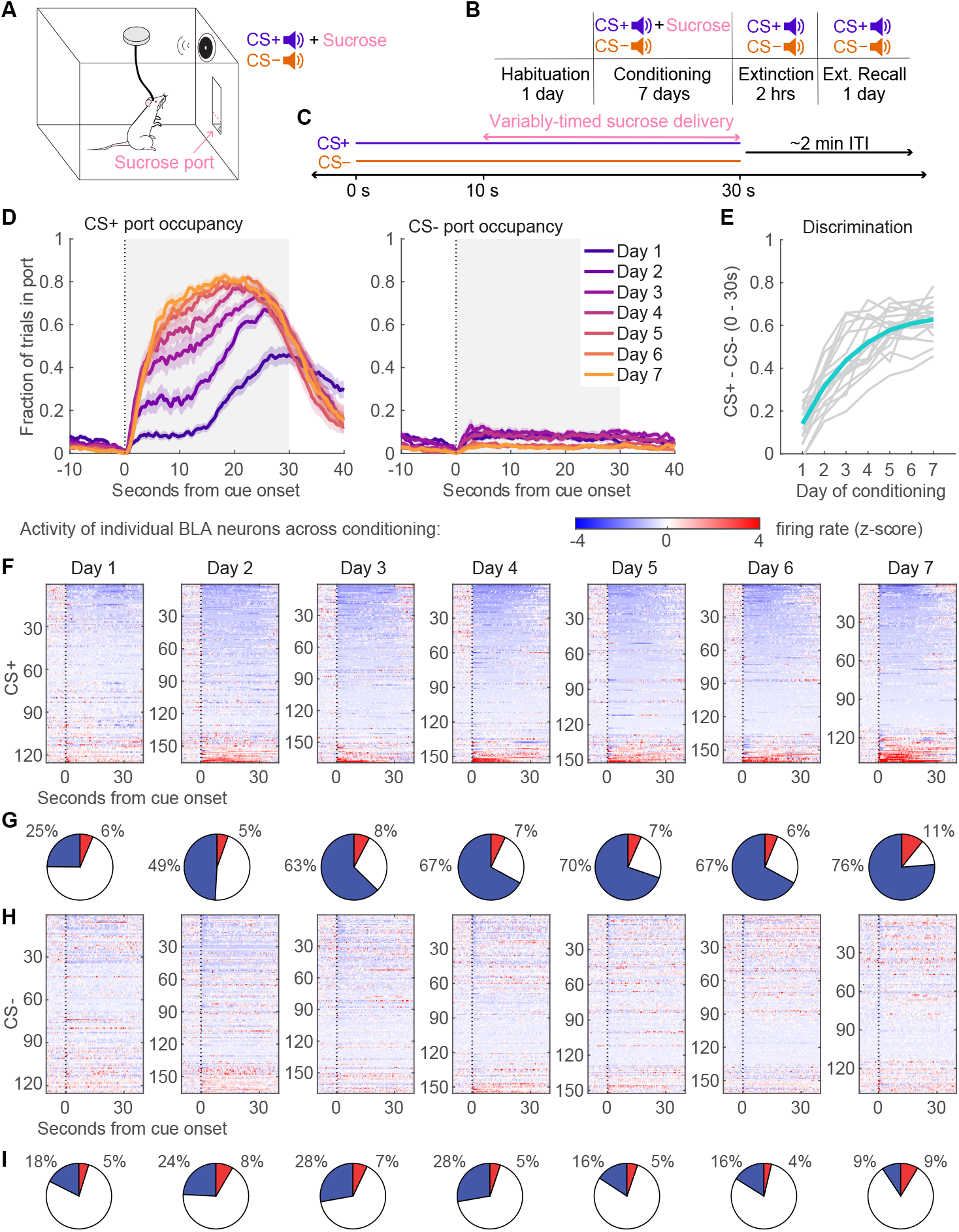
BLA activity during acquisition of conditioned responding to a CS+. (A) Behavioral apparatus. An auditory CS+ predicted delivery of sucrose into the reward port. (B) Rats (n=18) were conditioned for 7 days, followed by an immediate extinction session with no reward and an extinction recall the following day. (C) During Conditioning, reward was delivered at some point 10-24s following CS+ onset, except on rare trials (which were excluded from most analysis, see methods). (D) Fraction of time in sucrose port (mean ± SEM) on CS+ (left) and CS-(right) trials across Conditioning. (E) Difference between CS+ and CS-port occupancy during each 30s cue for individual rats (gray lines) and mean (blue) across Conditioning. (F) Z-scored, trial-averaged activity of individual BLA neurons aligned to CS+ across Conditioning. Neurons are sorted by their activity during CS+ (0 to 30s). (G) The proportion of BLA neurons with CS+-evoked inhibitions (blue) or excitations (red) across Conditioning. (H-I) As in (F-G), for CS-trials.

We recorded the activity of 125-169 (median = 155) neurons daily during Conditioning (Figure S1). Plotting the z-scored activity of all recorded neurons revealed an increasing prevalence of sustained activity during the CS+ across training (Figure 1F). The majority of cells tended to have reduced firing rate during the CS+ (around 70% in later sessions, Figure 1G). This was particularly striking given the low firing rates in BLA (mean = 1.5 Hz, median = 0.4 Hz) and the tendency for neurons with lower baseline firing rates to reduce their firing during CS+ (Figure S2). These sparse firing patterns encouraged us to analyze BLA activity at the population level. We implemented an approach that allowed us to track the activity of the entire neural population agnostic to the direction and statistical significance of individual neurons.

For each day of Conditioning, we found the population vector maximizing the difference between baseline (-15 to 0 sec) and initial CS+ (0 to 2 sec) firing. We then projected the activity of the population on CS+ and CS-trials onto this axis (the “CS+ dimension”), allowing us to assess the extent to which population activity resembled the initial CS+ response at different timepoints. First, we analyzed sustained cue-evoked activity, focusing on the time 2 to 10 sec following cue onset, which followed the phasic response we used to define the CS+ dimension and preceded reward delivery. Although population activity rapidly declined along the CS+ dimension following cue onset in initial sessions, we observed an increase in sustained CS+ activity on CS+ trials across Conditioning as well as increased discrimination between CS+ and CS-(Figure 2A-C). Notably, this sustained activity’s discrimination between CS+ and CS-correlated well with the rats’ behavioral discrimination of the two cues on a session-by-session basis (Figure 2D). We further looked at the emergence of the neural and behavioral responses on the first two days of Conditioning and found that this sustained activity was correlated with conditioned behavioral responding on a trial-by-trial basis for CS+ but not CS-(Figure 2E-G). These results indicate that behavioral conditioning is accompanied by sustained neural activity in BLA to the conditioned stimulus.

**Figure 2.**
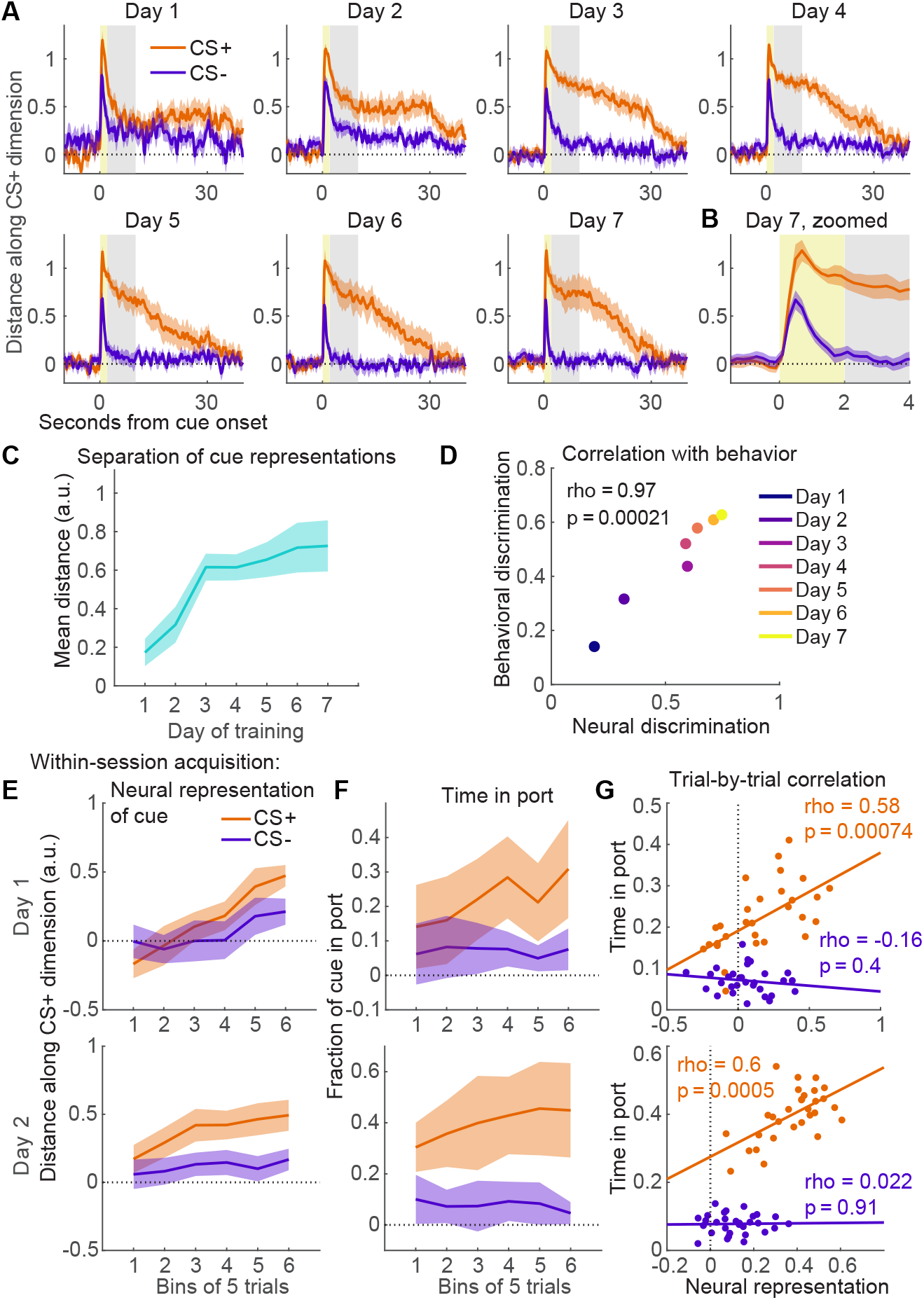
Sustained CS+ activity accompanies increases in conditioned responding. (A) BLA population activity along the CS+ dimension (defined as activity 0-2s from CS+ onset—yellow shading—relative to baseline activity -15-0s) on CS+ and CS-trials across the 7 days of Conditioning (mean ± SD, bootstrap). (B) Day 7 activity, zoomed in on cue onset. (C) Difference between CS+ and CS-activity 2-10s from cue onset (gray shading in (A)) (mean ± SD, bootstrap). (D) Correlation between behavioral (Figure 1E) and neural (C) discrimination of CS+ and CS-across Conditioning (Pearson’s rho = 0.97, p = 0.0002). (E) BLA population activity along CS+ dimension (defined as activity 0-2s from CS+ onset on the final 3 trials) 2-8s from cue onset, in bins of 5 trials for CS+ and CS-, on days 1 (top) and 2 (bottom) of Conditioning (mean ± SD, bootstrap). (F) Fraction of time in port during each cue for the first two days of Conditioning (mean ± SD). (G) Trial-by-trial time in port and distance along CS+ dimension for all 30 trials each of CS+ and CS-for first two days of Conditioning.

### Sustained CS+ activity tracks extinction of conditioned reward seeking

We next examined BLA activity during the extinction sessions. Conditioned behavioral responding decreased dramatically during Extinction and Extinction Recall (Figure 3A). If sustained BLA activity reflects conditioned responding, we would expect the BLA activity to decrease similarly across these sessions. The responses of individual BLA neurons were noticeably less pronounced during extinction (Figure 3C). Projecting population activity onto the CS+ dimension revealed decreased sustained activity on CS+ trials, resulting in poorer neural discrimination of the two cues (Figure 3D-E). Because the majority of conditioned behavior was extinguished during the first Extinction session, we further analyzed trial-by-trial changes in this session and found that the magnitude of sustained CS+ activity along the CS+ dimension and conditioned responding on CS+ trials were highly correlated (Figure 3F-G). Thus, sustained BLA activity is tightly linked to conditioned reward seeking in both acquisition and extinction of behavior.

**Figure 3.**
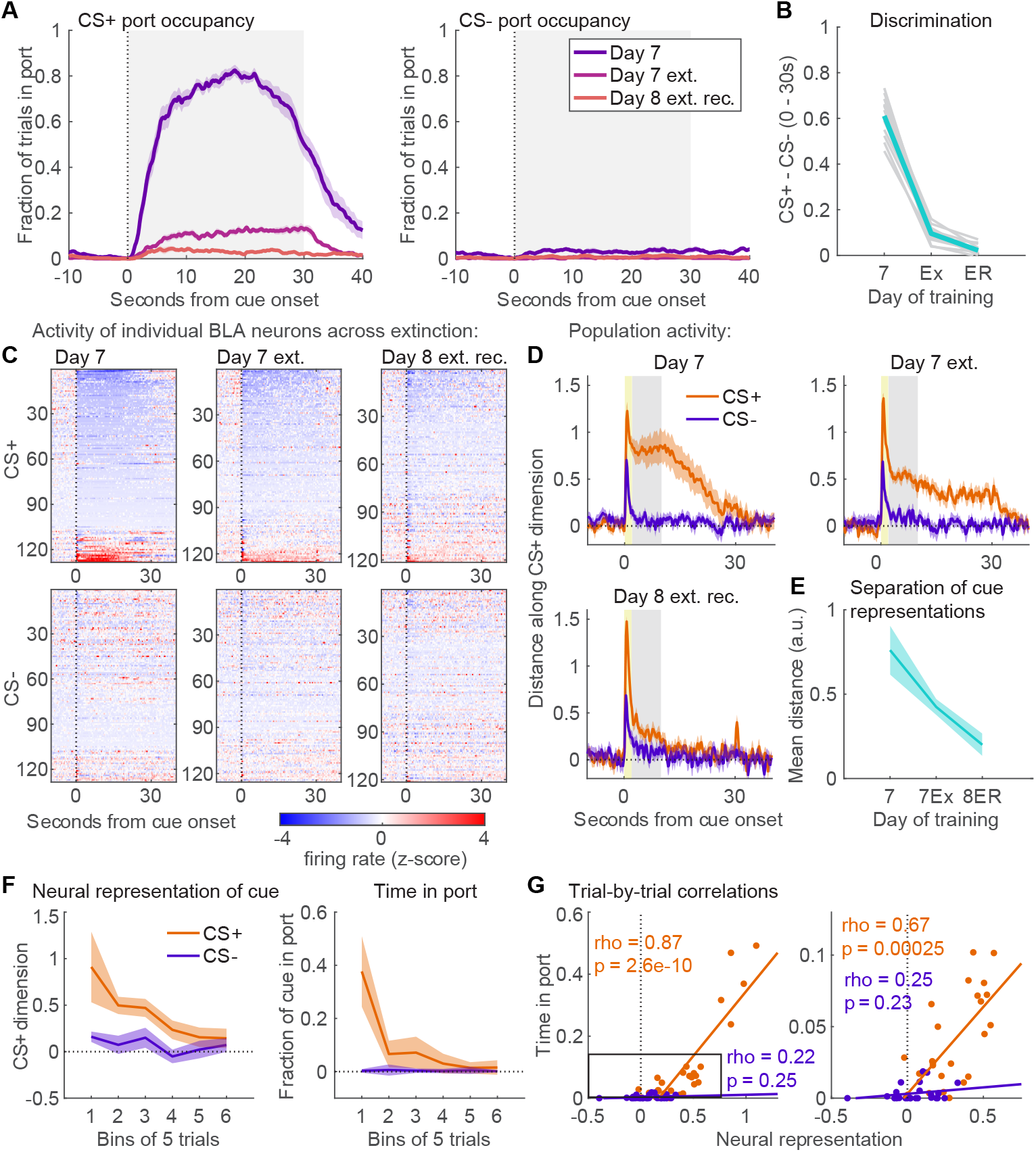
Sustained CS+ activity extinguishes along with behavior. (A) Fraction of time in sucrose port (mean ± SEM) on CS+ (left) and CS-(right) trials across Extinction. (B) Difference between CS+ and CS-port occupancy during each 30s cue. (C) Z-scored, averaged activity of individual BLA neurons aligned to CS+ (top) and CS-(bottom) across Extinction. Neurons are sorted by their activity 0-10s from CS+ onset. (D) BLA population activity along the CS+ dimension (defined as activity 0-2s from CS+ onset—yellow shading—relative to baseline activity -15-0s) on CS+ and CS-trials across Extinction (mean ± SD, bootstrap). (E) Difference between CS+ and CS-activity 2-10s from cue onset (gray shading in (D)) (mean ± SD, bootstrap). (F) Left: BLA population activity along CS+ dimension (defined as activity 0-2s from CS+ onset on the final 3 trials) 2-8s from cue onset, in bins of 5 trials for CS+ and CS-on first day of Extinction (mean ± SD, bootstrap). Right: Fraction of time in port (mean ± SD) for same session. (G) Trial-by-trial time in port and distance along CS+ dimension for all 30 trials each of CS+ and CS-for the first Extinction session.

### Sustained CS+ activity terminates with the conclusion of reward seeking

If sustained activity to the CS+ represents reward seeking, it follows that this sustained activity should terminate upon completion of reward-seeking behavior. To assess whether this was true, we analyzed BLA population activity along the CS+ dimension aligned to reward receipt (either reward delivery or when the rats first entered the port thereafter) and to the final port exit following reward consumption (Figure 4A). As sustained activity to the CS+ increased across Conditioning, we observed a corresponding decrease in activity following reward receipt (Figure 4B), resulting in significant decreases in activity along the CS+ dimension following reward on days 3 to 7 (p < 0.02, bootstrap; Figure 4C). These data indicate that the sustained BLA activity represents a reward-seeking state that initializes with cue onset and concludes with reward consumption.

**Figure 4.**
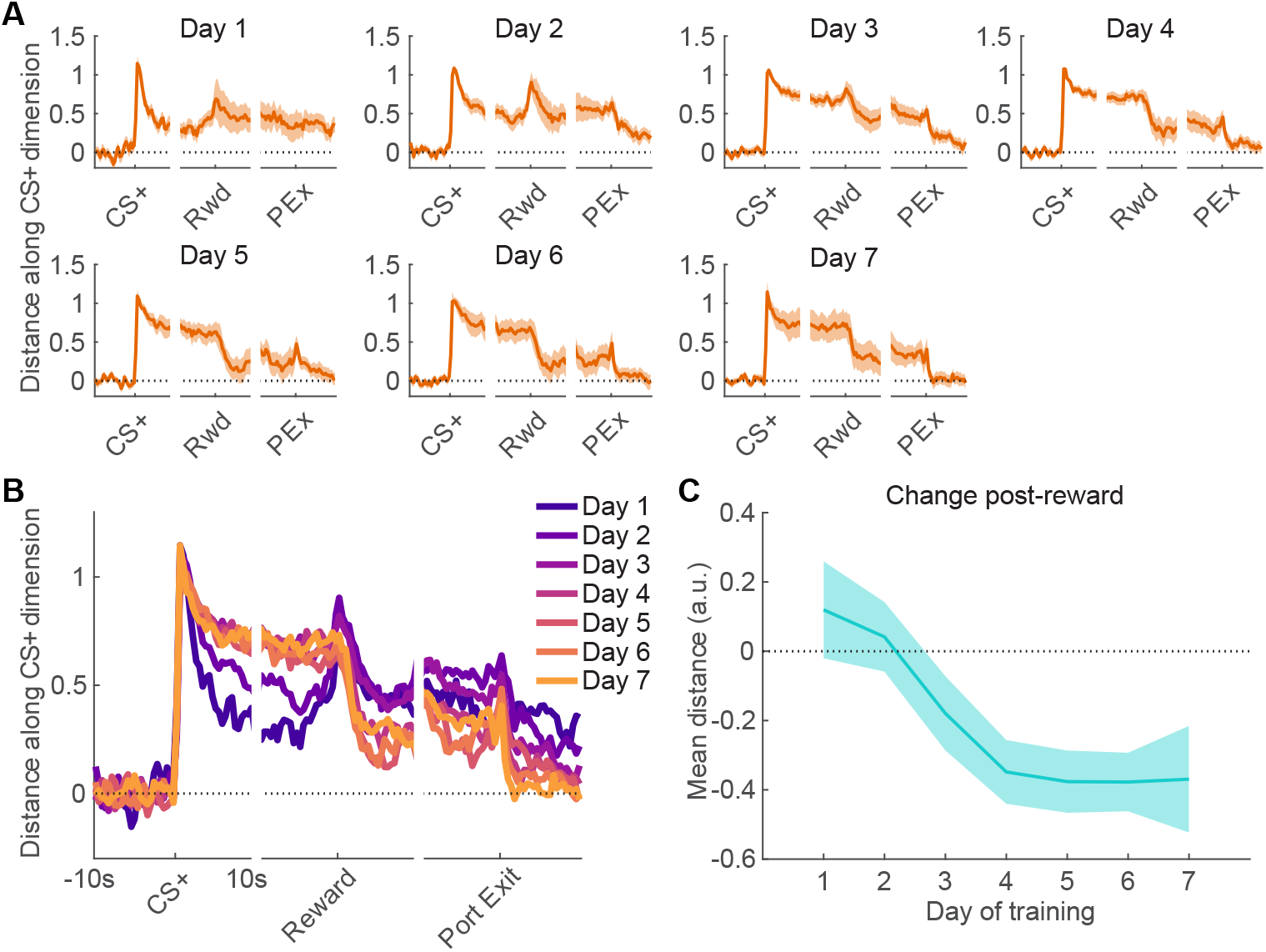
Sustained CS+ activity diminishes following reward receipt. (A) BLA population activity along the CS+ dimension (defined as activity 0-2s from CS+ onset relative to baseline activity -15-0s) on CS+ trials for the 10s before and after CS+ onset, reward delivery (or first port entry thereafter), and final port exit after reward consumption for the 7 days of Conditioning (mean ± SD, bootstrap). (B) The same plot as (A) without error bars to more easily compare sessions. (C) Difference in population activity along the CS+ dimension from 10s pre-reward to 10s post-reward for the 7 days of conditioning (mean ± SD, bootstrap).

### Disruption of sustained BLA activity compromises reward seeking

To determine if sustained BLA activity is necessary for reward seeking, we expressed channelrhodopsin (ChR2) in BLA neurons and implanted optical fibers directly above ChR2-expressing neurons (Figure 5A-B). Following Conditioning, we performed a test session where 50% of cue presentations were paired with 10s of laser-mediated stimulation of BLA neurons (Figure 5C). Because the majority of BLA cells had decreased firing rate during the CS+ (Figure 1F-G), this approach should disrupt normal population activity. We found that rats (n=9) spent significantly less time in the reward port during the first 10 seconds of the CS+ on stimulation trials relative to no laser trials (Figure 5D; p = 0.004, Wilcoxon signed-rank test), and this effect was stronger in ChR2 rats than YFP controls (Figure 5E; p = 0.005, Wilcoxon rank-sum test). These results are consistent with sustained BLA activity following the CS+ playing a causal role in conditioned reward seeking.

**Figure 5.**
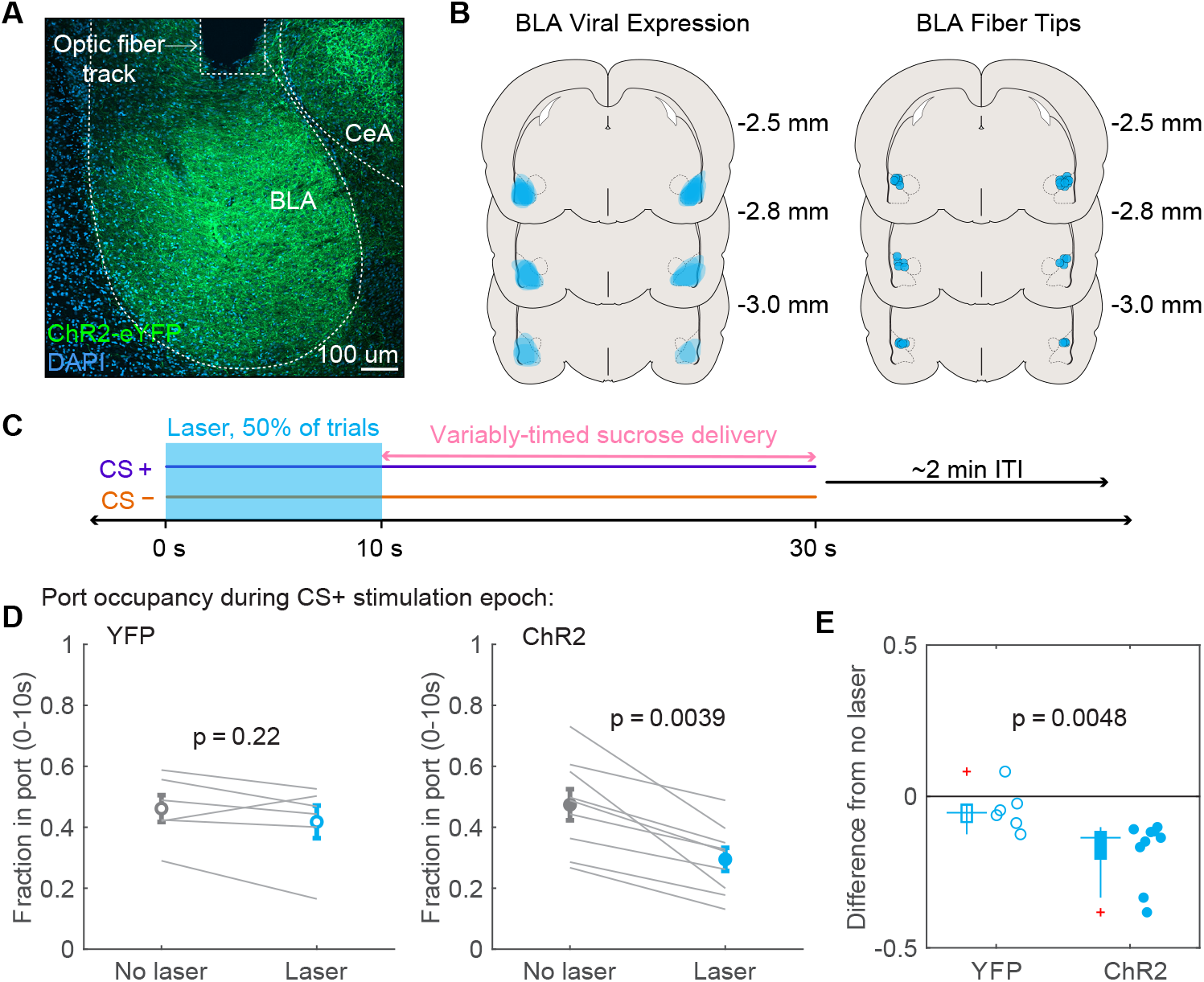
Sustained optogenetic excitation of BLA reduces conditioned reward seeking. (A) Example virus expression and optical fiber placement in BLA for optogenetic excitation. (B) Viral expression and fiber placement for rats included in the experimental group (n=9). (C) Experimental approach for optogenetic activation during cue-evoked behavior. Rats were trained on the Pavlovian discrimination described in Figure 1. On Day 8, blue light stimulation was paired with half of CS+ and CS-trials. (D) Fraction of time in port in the first 10 s of the CS+ during trials with (blue) and without (gray) light stimulation (mean ± SEM) and for individual rats (gray lines) (Wilcoxon signed-rank test). (E) Difference between laser and no laser trials for YFP and ChR2 groups (Wilcoxon rank-sum test).

### OFC is necessary for BLA control of conditioned reward seeking

The orbitofrontal cortex (OFC), which sends excitatory projections to BLA (Aggleton et al., 1980; Amaral and Insausti, 1992; Smith et al., 2000), is proposed to encode various aspects of the expected outcome during conditioned cues and convey this signal to BLA (Schoenbaum et al., 1998, 1999, 2011; Stalnaker et al., 2015). Thus, we tested the possibility that OFC drives the sustained CS+ activity we observed in BLA related to conditioned seeking of expected reward. To achieve this, we inactivated OFC via bilateral infusion of GABA agonists muscimol and baclofen (M/B) (or saline control) while recording individual BLA neurons (n=47 from saline sessions, n=55 from M/B) from a new group of rats (n=6) who underwent Conditioning (Figure 6A-B). Inactivation significantly reduced reward seeking during the CS+ compared to controls (Figure 6C-E). Accordingly, we found that, while BLA activity was unaffected by saline infusion, sustained activity to the CS+ along the CS+ dimension was significantly compromised following M/B infusion relative to controls (Figure 6F-G).

**Figure 6.**
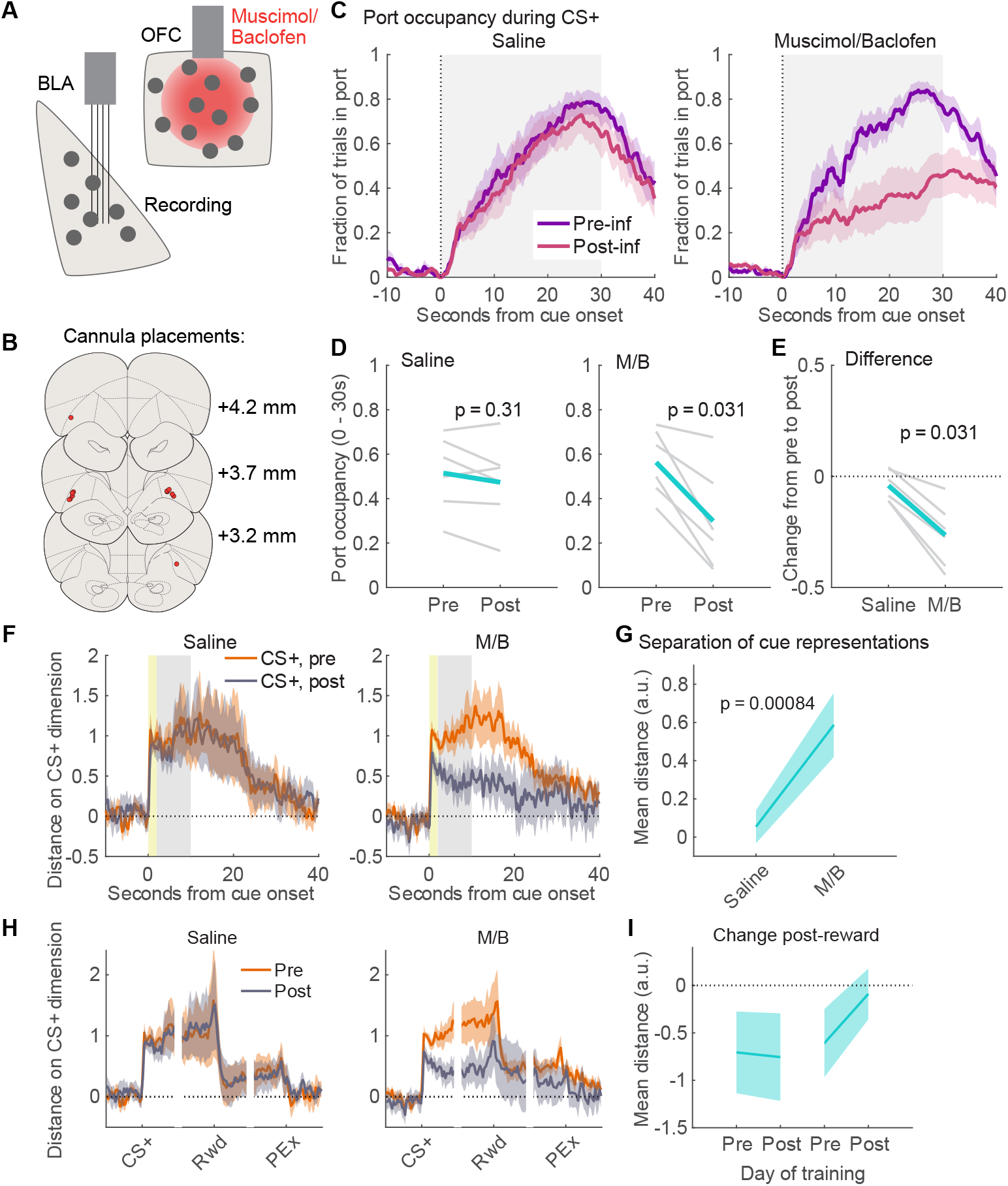
OFC contributes to BLA encoding of conditioned reward seeking. (A) Experimental approach to bilaterally inactivate OFC while recording in BLA. (B) Cannula placements in OFC (n=6). (C) Fraction of time in sucrose port (mean ± SEM) during CS+ pre- and post-infusion of saline (left) and muscimol/baclofen (M/B, right). (D) Change in CS+ port occupancy before and after infusion for saline (left) and M/B (right) for individual rats (gray) and mean (blue). Wilcoxon signed-rank test. (E) Difference in CS+ port occupancy for saline and M/B, individual rats (gray) and mean (blue). Wilcoxon rank-sum test. (F) BLA population activity along the CS+ dimension (defined as activity 0-2s from CS+ onset—yellow shading—relative to baseline activity -15-0s) on CS+ pre- and post-infusion for saline and M/B days (mean ± SD, bootstrap). (G) Difference between CS+ activity 2-10s from cue onset (gray shading in (F)) before and after infusion (mean ± SD, bootstrap). (H) BLA population activity along the CS+ dimension on CS+ trials for the 10s before and after CS+ onset, reward delivery (or first port entry thereafter), and final port exit after reward consumption for saline and M/B days (mean ± SD, bootstrap). (I) Difference between CS+ activity 10s before and 10s after reward delivery, before and after infusion (mean ± SD, bootstrap).

Moreover, while there was a pronounced decrease in activity along the CS+ dimension following reward receipt pre-infusion and post-saline (p < 0.01, bootstrap), this was not the case post-M/B (p = 0.36, bootstrap), although there was no significant change from pre-M/B (p = 0.09, bootstrap). Thus, inactivation of OFC appears to blunt sustained CS+ activity in BLA while simultaneously reducing conditioned reward seeking.

To more specifically test whether OFC input to BLA contributes to conditioned behavior, we expressed the light-sensitive inhibitory opsin ArchT3.0 in the OFC and placed optical fibers in the BLA to allow selective inhibition of OFC terminals in the BLA upon delivery of green laser (Figure 7A-B). We performed the same training and test as the ChR2 group (Figure 5; Figure 7C). On the test day, inhibition of OFC terminals in BLA (n=10) during the initial 10 sec of CS+ significantly reduced the time spent in the port compared to CS+ trials with no inhibition (Figure 7D; p=0.006, Wilcoxon signed-rank test), and this effect was greater in ArchT3.0 rats than YFP controls (Figure 7E; p = 0.03, Wilcoxon rank-sum test). These findings show that a direct projection from OFC to BLA contributes to conditioned reward seeking. Taken together, the OFC manipulations suggest that OFC input is critical for BLA sustained activity to the CS+, which drives conditioned reward seeking.

**Figure 7.**
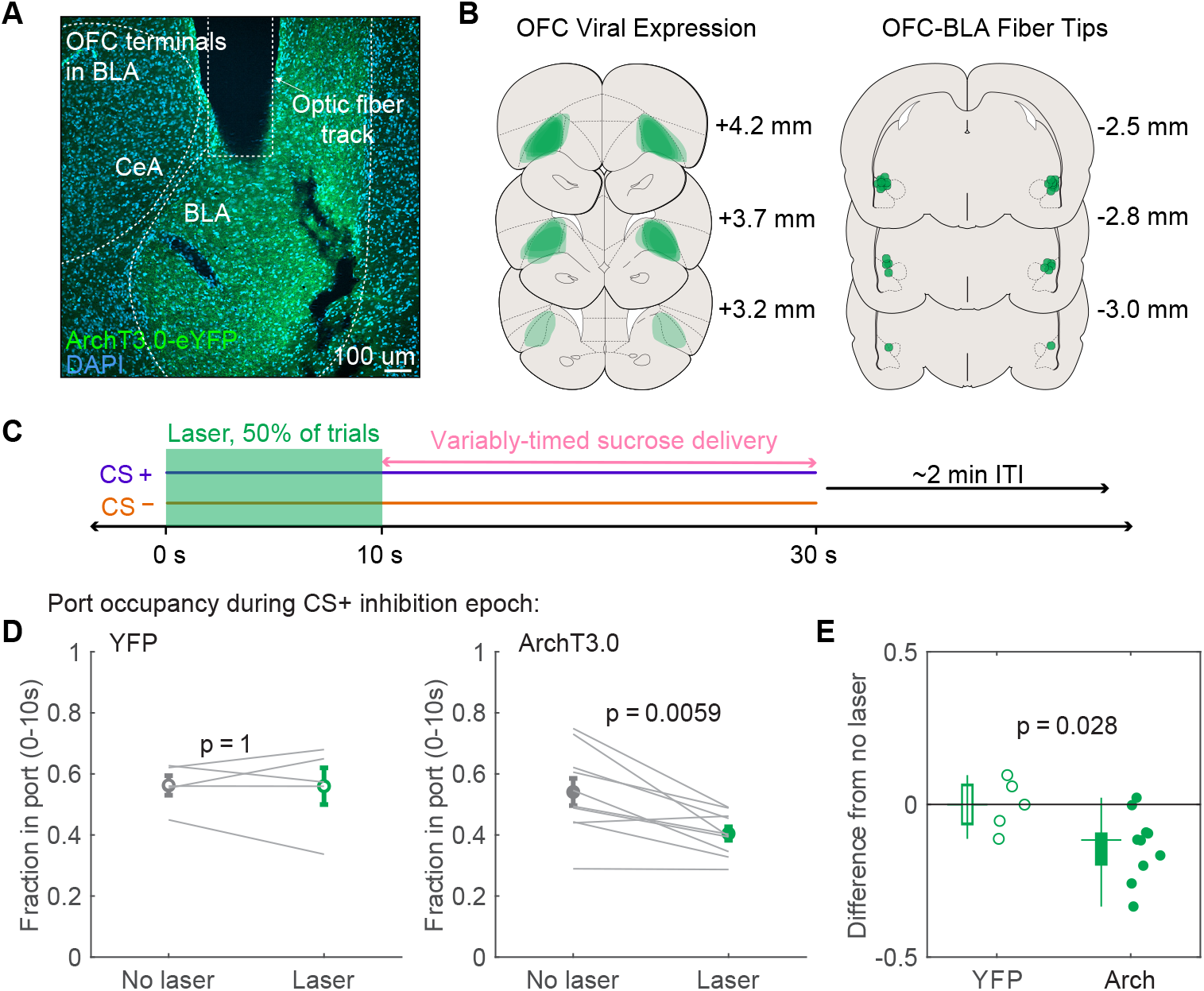
Sustained optogenetic inhibition of OFC inputs to BLA reduces conditioned reward seeking. (A) Example virus expression in OFC terminals and optical fiber placement in BLA for optogenetic inhibition. (B) Viral expression and fiber placement for rats included in the experimental group (n=10). (C) Experimental approach for optogenetic inhibition during cue-evoked behavior. Rats were trained on the Pavlovian discrimination described in Figure 1. On Day 8, green light inhibition was paired with half of CS+ and CS-trials. (D) Fraction of time in port in the first 10 s of the CS+ during trials with (green) and without (gray) light stimulation (mean ± SEM) and for individual rats (gray lines) (Wilcoxon signed-rank test). (E) Difference between laser and no laser trials for YFP and ArchT3.0 groups (Wilcoxon rank-sum test).

## DISCUSSION

Our study provides new data indicating that BLA neuronal population activity encodes a reward-seeking state during appetitive Pavlovian conditioning. This neural activity, primarily consisting of reductions in firing below baseline levels, developed over the course of task acquisition and rapidly diminished during extinction. This sustained neural activity change closely correlated with the quantity of reward seeking behavior and, compellingly, terminated upon acquisition of reward, indicating a tight link between BLA neural activity and the reward-seeking component of the behavior. The importance of this activity for reward seeking was further demonstrated with optogenetic activation of BLA, which reduced conditioned responding during the CS+. We also found an important role for OFC in the contribution of BLA activity to this task; inactivation of OFC disrupted the sustained CS+-evoked neural responses in BLA, and optogenetic inhibition of OFC axon terminals in the BLA reduced reward seeking. Our study provides new insight into how OFC-dependent BLA neural activity orchestrates ongoing behavior during conditioned reward seeking.

BLA is known to be a major site of plasticity underlying cue-outcome learning (Fanselow and LeDoux, 1999; Janak and Tye, 2015). After conditioning, thalamic inputs onto lateral amygdala are strengthened (McKernan and Shinnick-Gallagher, 1997; Tye et al., 2008; Clem and Huganir, 2010) and cue-evoked activation of lateral amygdala neurons is potentiated (Quirk et al., 1995; Rogan et al., 1997; Tye et al., 2008). This enhanced cue-evoked LA activity is proposed to promote conditioned behavioral responses through its connectivity with basal amygdala and further downstream regions (LeDoux et al., 1990; Maren and Quirk, 2004; Duvarci and Pare, 2014; Janak and Tye, 2015; Namburi et al., 2015). In the simplest version of this model, BLA cue-evoked excitations are transferred to the CeA to produce freezing while for appetitive conditioning, and BLA cue-evoked excitations are transferred to the NAc (Ambroggi et al., 2008; Duvarci and Pare, 2014; Janak and Tye, 2015; Namburi et al., 2015; Tovote et al., 2015). This simplistic model does not capture the observed heterogeneity in BLA cue-evoked activity (excitations and inhibitions for appetitive and aversive cues) among subpopulations sending axons to NAc, CeA, and ventral hippocampus (Beyeler et al., 2016). In our study, we found that the majority of cue-evoked responses were decreases in firing, which is consistent with data from other tasks with extended cue periods in freely moving rats (Lee et al., 2017; Kyriazi et al., 2018). The repeated demonstration of CS+-evoked inhibitions, as well as our finding that optogenetic activation of the BLA disrupts reward seeking (rather than promoting it), emphasizes the importance of integrating BLA inhibitions into circuit models of conditioned reward seeking.

In our work, we found more evidence of the important role the OFC plays as an input to the BLA. The OFC is known to contribute to BLA neuronal responses to cues, particularly in settings where cue-outcome associations are changing (Saddoris et al., 2005; Morrison et al., 2011; Lucantonio et al., 2015; Sharpe and Schoenbaum, 2016; Arguello et al., 2017). We found that the OFC is critical for the persistent population activity we observed in BLA during reward seeking and, specifically, that OFC input to the BLA contributes to conditioned behavior in this task. Interestingly, multiple previous reports find that simple Pavlovian behavior, such as port entry in response to a reward-predictive cue, is generally unimpaired by either OFC or BLA manipulation; rather, the OFC and BLA roles are restricted to occasions in which a cue-evoked representation of the reward, including its specific features and expected value, is required for appropriate behavior (Hatfield et al., 1996; Gallagher et al., 1999; Pickens et al., 2003, 2005; Ostlund and Balleine, 2007; Burke et al., 2009; Takahashi et al., 2009; Fraser and Janak, 2023). Similarly, electrophysiological and pharmacological studies have suggested that interactions between the OFC and the BLA are required for cue-dependent behavior when information about the expected value is needed (Baxter et al., 2000; Schoenbaum et al., 2003; Izquierdo and Murray, 2004; Saddoris et al., 2005; Churchwell et al., 2009; Lucantonio et al., 2015; Sias et al., 2021). In contrast, we found Pavlovian conditioned behavior was sensitive to OFC and BLA manipulations, and we speculate that it is due to the increased task demands introduced by an unpredictable reward delivery time and the prolonged cue time, which may recruit OFC-dependent sustained activity in BLA more prominently.

A particularly interesting feature of our results is the cue-driven emergence of a neural state that persists until the acquisition of reward. This suggests that, in this Pavlovian task, BLA activity encodes conditioned reward seeking. These data are consistent with other findings that BLA activity is closely linked to reward seeking states. For instance, the magnitude of cue-evoked neural modulation depends on whether subjects execute reward seeking behavior (Lee et al., 2016), and the activity during CS+ presentation and reward-related behaviors is highly correlated in rewarding contexts (Kyriazi et al., 2018). The idea that BLA neural activity represents state encoding extends beyond simple Pavlovian conditioning. During self-paced reward seeking, BLA activity differentiates between reward seeking and reward consumption behavioral phases (Courtin et al., 2022). Moreover, BLA activity, and OFC input to BLA, are necessary for learning reward-specific states, further supporting the notion that BLA activity plays a role not just in promoting ongoing behavior but also in learning from that experience (Sias et al., 2021).

In conclusion, our data demonstrate the importance of sustained population dynamics in BLA, and the contribution of OFC input, for the execution of conditioned reward seeking. These results highlight the continued importance of studying the role of BLA in state coding, especially given the purported involvement of BLA in aberrant reward seeking conditions, including addiction (Wassum and Izquierdo, 2015).

## METHODS

### Subjects

Adult male Sprague Dawley rats (Harlan, Indianapolis, IN) had access to ad libitum food and water until 2 days prior to the onset of behavioral training and neural recording at which time they were restricted to 14.5-16.5g of food per day, maintaining ∼90% of pre-restriction body weight. All procedures were approved by the Institutional Animal Care and Use Committees of UCSF, the Ernest Gallo Clinic and Research Center, and Johns Hopkins University, in accordance with guidelines of the US National Institutes of Health.

### Surgery

For BLA neural recording, fixed microelectrode arrays (NeuroBiological Laboratories, Denison, TX) were implanted bilaterally in the BLA (2.8 mm posterior to bregma; +4.6 mm lateral to midline; 7.0 mm ventral to dura). For pharmacological inactivation, in addition to BLA electrode arrays, bilateral 26-gauge stainless steel guide cannulae (Plastics One, Roanoke, VA) were implanted aimed at OFC (3.5 mm anterior to bregma; 2.8 mm lateral to midline; 2.7 mm ventral to skull). Guide cannulae were targeted 2 mm above the intended infusion site because infusion cannulae extended this distance beyond the end of the guide. For BLA photoactivation, rats were infused in the BLA bilaterally with adeno-associated viral vectors, AAV5-CaMKII-ChR2-YFP (ChR2; UNC Vector Core, Chapel Hill, NC) or AAV5-CaMKII-YFP (YFP; UNC Vector Core), to express channelrhodopsin-2 (ChR2)-YFP or YFP alone (0.5μl/ side at 0.1μl/min; coordinates above) and 300-μm optic fibers were implanted 0.1 mm above injection sites. For OFC axon terminal photoinhibition, rats were infused bilaterally with AAV5-CaMKII-ArchT3.0-YFP (UNC Vector Core) or the YFP control virus into the OFC (coordinates above), and implanted bilaterally with optic fibers above the BLA, as above.

### Pavlovian discrimination conditioning

Three days before the start of behavioral training, rats were pre-exposed to sucrose in the home cage for 48 hours. Rats were then trained on a Pavlovian discrimination procedure for ∼2 hrs/day for 7 days. Conditioning consisted of 30 presentations each of two auditory stimuli, a 2.9 kHz tone or white noise, delivered for 30s, at ∼70dB, with one stimulus paired with delivery of 0.2ml of a 10% sucrose solution over 6 s. The ITI was a 2 min variable interval, although if the rat was in the reward port at ITI end, 10s was added to the ITI until the rat was not in the port at ITI end. The stimulus paired with sucrose reward was counterbalanced across animals. Sucrose delivery during the cue was variable, occurring with equal probability at each integer value between 10 and 24 s after cue onset for ∼94% of trials, but occurring after 1s on ∼6% of trials. These trials were excluded from analysis of behavior and neural activity (except for the trial-by-trial correlations in Figure 2F-G and Figure 3F-G). Sucrose delivery latency was variable to discourage conditioned responding based on timing and to encourage conditioned responding throughout the cue period.

### Extinction training and testing

Extinction training was conducted on Day 7 of Pavlovian discrimination training: the 7th conditioning session was modified to include 60 presentations each of the CS+ and CS-, and after the first 30 CS+ presentations, the pump drive was manually retracted from the end of the syringe containing sucrose, such that the last 30 CS+ presentations were unreinforced. On the following day, rats underwent a second extinction session consisting of 30 unreinforced presentations of both the CS+ and CS- to test for extinction learning recall.

### Pharmacological inactivation of OFC

Rats received 0.5 ml infusions (0.3 ml/min) of a cocktail (M/B) of the GABAA agonist muscimol (250ng/ml) and the GABAB agonist baclofen (250ng/ml) or saline as described (Keiflin et al., 2013). Infusion cannulae were left in place for 2 min after infusion to allow for diffusion, and subjects returned to the conditioning chamber after ∼8 min.

### Optogenetic activation of BLA

Rats infused with an AAV5-CaMKII-ChR2-YFP or AAV5-CaMKII-YFP and fiber optic implants in the BLA received the 7-day Pavlovian discrimination procedure as described above, except that the pseudorandom ITI was adhered to whether or not the subject was in the reward port. During this period, rats were tethered with an optic cable to a liquid swivel rotary joint, which was connected to a 100mW 473nm blue DPSS laser (OEM Laser Systems, Midvale, UT), but no light was delivered. On day 8, light (5-ms pulses, 20 Hz) was delivered during the first 10 s of 50% of CS+ and 50% of CS-trials. Sucrose was available as usual.

### Optogenetic inhibition of OFC terminals in BLA

Rats infused with AAV5-CaMKII-ArchT3.0-YFP or AAV5-CaMKII-YFP in OFC and fiber optic implants in the BLA were tethered to a 532nm green DPSS laser and trained as in the BLA optogenetic activation study. On test day, light (10-s constant pulse) was delivered during the first 10 s of 50% of CS+ and 50% of CS-trials, and sucrose was available as usual.

### Analysis of neural activity

To detect excitations and inhibitions in response to the CS+ and the CS-, we compared the number of spikes in the 10s following each cue presentation to the 10s preceding each cue presentation and performed a Wilcoxon signed-rank test, with a cutoff of p < 0.05.

PSTHs were created by binning the neural activity into 20ms bins surrounding the events of interest. These PSTHs were first smoothed on a per trial basis with half normal filter with sigma = 5 so activity was only smoothed over previous activity (no influence of upcoming activity on the trace). The trial-averaged PSTH was then smoothed with another half normal filter with sigma = 15. These PSTHs were then z-scored by subtracting the mean baseline activity (before all cue onsets) and dividing by the standard deviation (across all trials).

To project population activity onto the CS+ dimension (Ottenheimer et al., 2023), we normalized the unsmoothed firing rate of each neuron in a PSTH spanning baseline and CS+ activity so that each neuron’s activity ranged from 0 to 1. We then calculated the difference between each neuron’s CS+ evoked activity (0 to 2s) and baseline activity (-15 to 0s), producing a vector describing how modulated each neuron is to the CS+ relative to its total range of firing. We then projected population activity onto this axis by finding the the dot product of this vector with the normalized activity (0 to 1, as before) at baseline (-15 to 0s), CS+ onset (0 to 2s), sustained CS+ and CS-responses (2 to 8s), and smoothed PSTHs surrounding CS+, CS-, reward receipt, and final port exit following reward consumption. All of these population projections were linearly transformed such that 0 was the projected population activity at baseline and 1 was the projected population activity as CS+ onset. The resulting value is essentially a weighted average of population activity where the weights are determined by how modulated each neuron is by the CS+ onset.

Because this method could potentially be dominated by a few neurons with very large CS+ onset modulations, we ran control analyses. First, instead of reporting the singular population projection, we report the mean ± SD of bootstrapped projections, with which, by selecting 10000 groups of neurons (with replacement), we avoid reliance on a particular combination of neurons. We also bootstrapped the difference between CS+ and CS-sustained activity (2 to 8s) and pre- and post-reward receipt (10s each), allowing derivation of p values. Second, we reran our analysis, excluding neurons in the top 10% of CS+ modulations, and found the same pattern of results (all bootstrap tests and correlations still significant at p < 0.05 except for reduction along CS+ dimension post-reward on Conditioning day 3).

To look at trial-by-trial projected population activity across acquisition and extinction, we used the same approach, but with the CS+ evoked activity (0 to 2s) in the final (acquisition) or first (extinction) three trials to define the axis, and then projected the average sustained CS+ and CS-activity (2 to 8s) on each of the 30 trials onto the axis. We used Pearson correlation coefficients to relate this activity to trial-by-trial port entry behavior.

Our analysis was conducted on all recorded neurons. To ensure that the results were robust among principal cells and not driven by putative interneurons, we ran the same analysis, excluding all neurons with baseline firing rate greater than 6 Hz (Kyriazi et al., 2018) (making up 5.8% of the recorded population), and found the same pattern of results.

### Behavioral measures and analysis

The fraction of time rats spent in the sucrose port during CS presentation (port occupancy) was calculated by dividing the amount of time the rat’s nose was in the port during the CS by the total duration (30s). Trials in which the reward was delivered 1s after CS+ onset (∼6%) were excluded from analysis (except for the trial-by-trial correlations in Figure 2F-G and Figure 3F-G). For the optogenetic experiments, we performed the same analysis on the first 10s (during laser stimulation). Reward receipt was defined as the first moment the rat was in the port following reward delivery. Final port exit following reward was defined as the first port exit following reward delivery with more than 3 seconds until the next port entry.

Wilcoxon signed-rank tests were used to determine whether % time in port changed before or after OFC pharmacological inactivation, and on trials with and without laser stimulation for optogenetic manipulations. The amount of change in port entry time was compared across groups with Wilcoxon rank-sum tests.

## Funding and Acknowledgements

This work was supported by US National Institutes of Health grants R01 DA036996, R01 DA035943, R01 AA027213, F32 DA036996, R00 DA042895, R01 MH129370, and R01 MH129320, and funds from the State of California for medical research on alcohol and substance abuse through UCSF. We wish to acknowledge Anthony Mefford, Rebecca Reese, Lacey Sahuque, and Chan Hyouk Park for technical contributions to these experiments.

## Supplemental Figures

**Figure S1.**
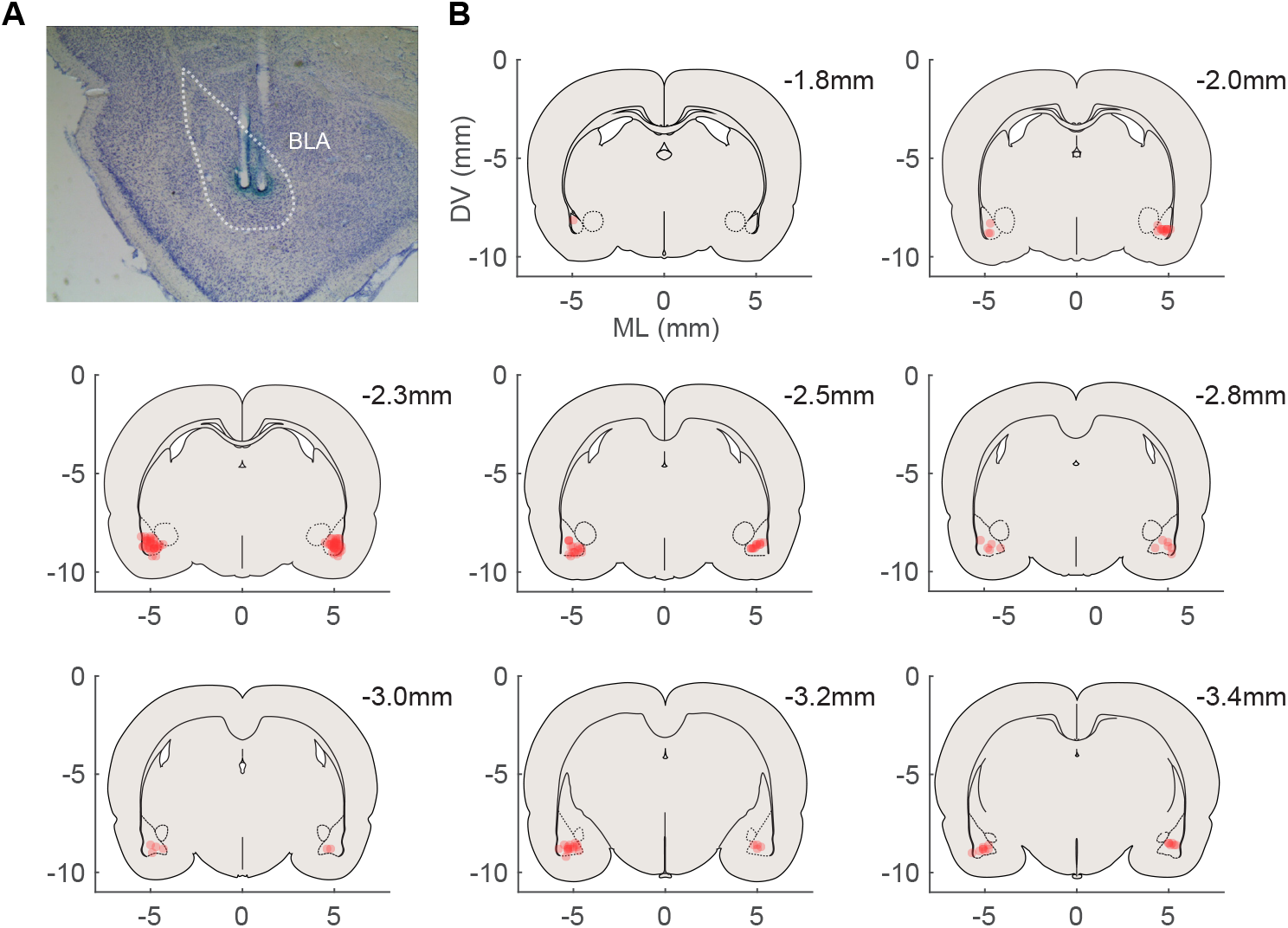
Placements for electrophysiology. (A) Histology image showing example electrode placement. (B) Placements of individual wires included in the Conditioning and Extinction datasets. Placements for the OFC inactivation experiment followed the same coordinates and inclusion criteria.

**Figure S2.**
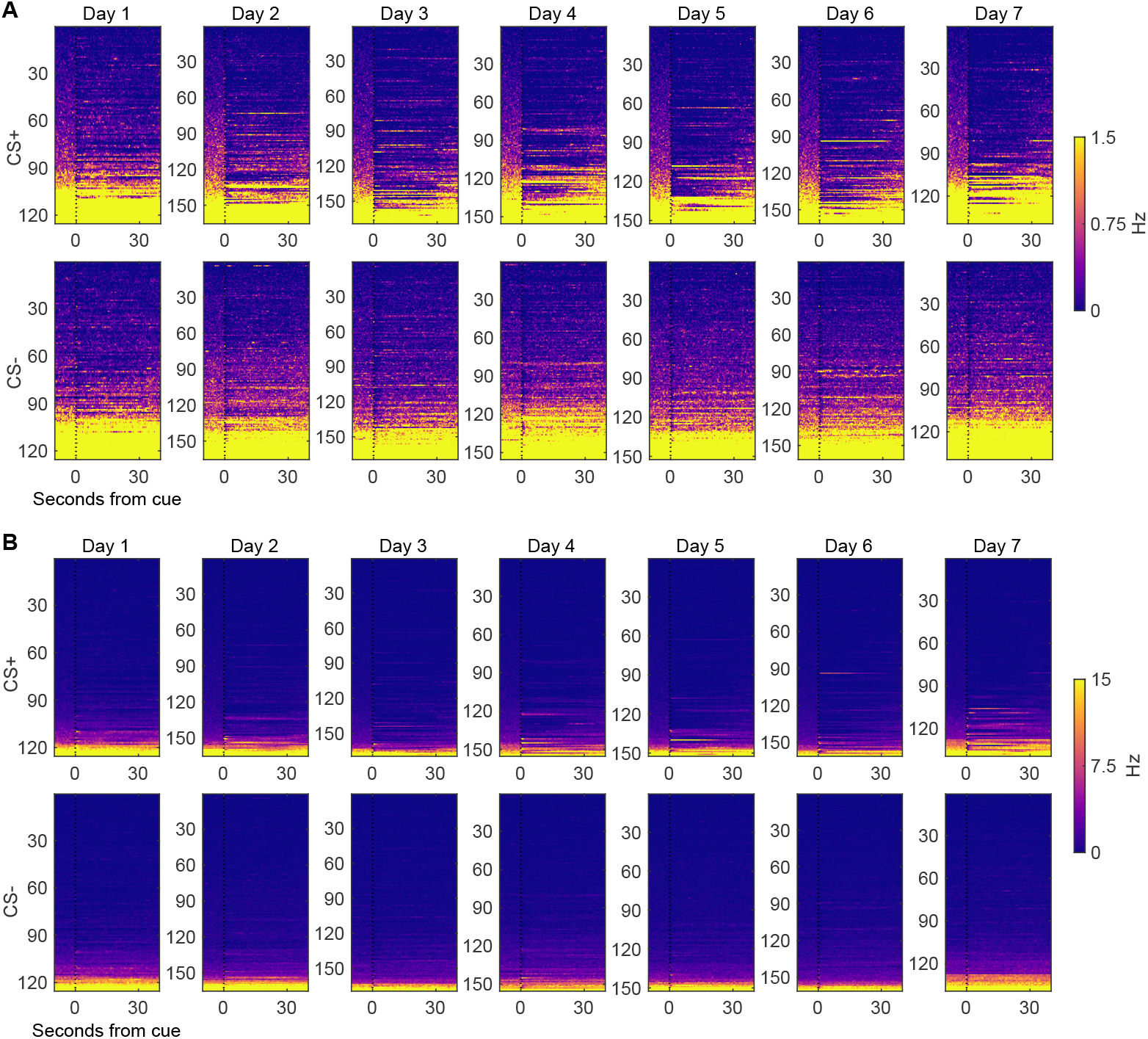
Unnormalized activity of neurons recorded during Conditioning. (A) Activity of all recorded BLA neurons during conditioning, sorted by their baseline firing rate, on CS+ (top) and CS-(bottom) trials. (B) The same activity, recolored to allow better visualization of neurons with higher firing rates.

